# Exploring the structural diversity and evolution of the D1 subunit of photosystem II using AlphaFold and Foldtree

**DOI:** 10.1101/2025.03.06.641835

**Authors:** Tom Dongmin Kim, Daniella Pretorius, James W. Murray, Tanai Cardona

**Affiliations:** School of Biological and Behavioural Sciences, Queen Mary University of London, United Kingdom; Department of Life Sciences, Imperial College London, United Kingdom

## Abstract

While our knowledge of photosystem II has expanded to time-resolved atomic details, the diversity of experimental structures of the enzyme remains limited. Recent advances in protein structure prediction with AlphaFold offer a promising approach to fill this gap in structural diversity in non-model systems. This study used AlphaFold to predict the structures of the D1 protein, the core subunit of photosystem II, across a broad range of photosynthetic organisms. The prediction produced high-confidence structures, and structural alignment analyses highlighted conserved regions across the different D1 groups, which were in line with high pLDDT scoring regions. In contrast, varying pLDDT in the DE loop and terminal regions appear to correlate with different degrees of structural flexibility or disorder. Subsequent structural phylogenetic analysis provided a phylogeny that is in good agreement with previous sequence-based studies. Moreover, the phylogeny supports a parsimonious scenario in which far-red D1 and D1^INT^ evolved from the ancestral form of G4 D1. This study demonstrates the potential of AlphaFold in studies on structural diversity and the evolution of photosynthesis.

## Introduction

Oxygenic photosynthesis is one of nature’s most remarkable innovations, providing a foundation for life by converting light energy into chemical energy at a global scale. The process begins with Photosystem II (PSII), where photons drive the oxidation of water to electrons, protons, and molecular oxygen. Elucidating the structure, function, and evolution of PSII has been one of the key aims of photosynthesis research, and advancements in structural biology over the previous decades have led to a highly detailed atomic-level understanding of the enzyme. Not only in terms of the mechanism of water oxidation, but also in terms of the assembly and repair process and on matters of origin and evolutionary adaptations (Oliver et al., 2023, Gisriel, 2024, Komenda et al., 2024, Yano et al., 2024). As of January 2025, the Protein Data Bank (Berman et al., 2000) contains 140 experimentally determined structures of PSII. However, while progress has been made in resolving structures from an increasingly expanding diversity, these structures still represent a limited range of photosynthetic organisms. Of the 140 entries, 105 originate from *Thermosynechococcus vestitus* (*T. elogatus*) or *Thermostichus vulcanus* (*Thermosynechococcus vulcanus*), and a further 12 are from *Synechocystis* sp. PCC 6803 or *Chlamydomonas reinhardtii*, accounting for nearly 84% of the dataset. The remaining 23 structures belong to other cyanobacteria, and a range of eukaryotic algae and plants.

Recent developments in protein structure prediction have transformed the field, with AlphaFold (AF) leading the way (Jumper et al., 2021). By achieving unprecedented accuracy in protein structure prediction, AlphaFold has provided researchers with an alternative approach to explore protein structures in cases where experimental data are unavailable or challenging to obtain. Its utility has been demonstrated in a wide range of applications, including experimental structure determination (Terwilliger et al., 2023), elucidating protein-protein interactions (Yu et al., 2023), and in novel protein design (Wicky et al., 2022).

The D1 subunit of PSII is a key component of the enzyme, providing most of the ligands to the oxygen evolving Mn_4_CaO_5_ cluster (Umena et al., 2011) and controlling many of the photochemical properties of the enzyme (Cardona et al., 2012, Rutherford et al., 2012). Within cyanobacteria, a large diversity of D1 subunits evolved via a gene duplication process that started before the extant lineages radiated and which has never stopped, see Oliver et al. (2023) for a recent detailed review. The evolution of D1 isoforms, not only enables cyanobacteria strains to adapt to varying conditions such as high light, low light, or low oxygen, but it also has enabled the enzyme to acquire novel functions, such as chlorophyll *f* synthesis.

Here, we explored the extent to which AlphaFold-predicted structures can be used to extract novel insights into the diversification of PSII, focusing on the reaction centre subunit D1. We also explored the extent to which phylogenetic trees inferred with these predicted structures can be used to inform photosystem evolution.

## Materials and Methods

### Structural prediction with AlphaFold

A total of 538 cyanobacterial D1 amino acid (AA) sequences were downloaded using BLAST for a previous study (Saw et al., 2021). These sequences included a representative sampling of all known groups, curated at a redundancy level of 98% sequence identity using a custom-made python script (see Data Availability). A further 293 sequences were collected from photosynthetic eukaryotes (Archaeplastida) on 27 April 2021, using BLAST, not including algae with secondary plastids, and sampled down to 98% sequence identity. A subset of 98 cyanobacterial D2 sequences were also included in the analysis to be used as an outgroup in phylogenetic analysis.

Structural prediction of the AA sequence set was done with Colabfold (Mirdita et al., 2022), installed locally in high performance computing cluster (HPC) at Imperial College London (see DOI: 10.14469/hpc/2232) and Aprocrita HPC at Queen Mary University of London (DOI: 10.5281/zenodo.438045). Colabfold is an accelerated version of AlphaFold2 using MMseq2 (Steinegger and Soding, 2017) for Multiple Sequence Alignment (MSA) generation. It generates 5 predicted structures per input sequence and ranks them by average pLDDT score. The highest ranking model was then taken forward for further analysis.

### Structural alignment analysis with representative structures

For visual analysis of the predicted structures from different D1 groups, three structures were selected to represent each of the following groups, using the categorisation introduced by Cardona et al. (2015) and Sheridan et al. (2020): G1, G2, G3, G4 in Gloeobacterales (G4*Glo*), the standard D1 in the far-red light photoacclimation response (FaRLiP) gene cluster (D1^FRL^), standard G4 in Cyanobacteria (G4Bac) and G4 in Eukaryotes (G4Euk). In the case of G0, there are only two sequences known, from *Gloeobacter kilaueensis* and *Gloeobacter morelensis*, and both were included (Saw et al., 2021). G3 and G4 represent the standard form of D1 capable of sustaining water oxidation if assembled into a PSII complex, and G0-G2 represent the atypical forms of D1 characterised by lacking some of the ligands to the Mn_4_CaO_5_ cluster (Murray, 2012, Cardona et al., 2015). The complete list of D1 is in Table S1 in supplementary material. The 24 AF structures were aligned by mTM-Align (Dong et al., 2018) with default settings, together with a D1 extracted from the CryoEM structure from *Synechocystis* sp. PCC 6803 by Gisriel et al. (2022b), PDB 7N8O, as a reference. The program provides a matrix of Root Mean Square Distance (RMSD) and TM score from all-to-all alignment results, as well as the common core of aligned structures defined as conserved domain structures without gaps and within 4 Å of maximum pairwise distance. Molecular graphics and analyses were performed with UCSF ChimeraX (Meng et al., 2023).

### Structural phylogenetics of D1 AF structures

All AF structures were fed to Foldtree (Moi et al., 2023) to create a phylogenetic tree using default settings. In short, Foldtree is a python based workflow made with Snakemake (Koster and Rahmann, 2012), starting from all-to-all structural alignment with Foldseek (Van Kempen et al., 2024). Foldseek compresses the local structural context between closest neighbouring C_α_s into a structural alphabet (3Di), using a pre-trained Vector quantised variational autoencoder (van den Oord et al., 2017), enabling fast alignment of the new structure-aware sequences. It combines AA and 3Di substitution scores with weights 1.4 and 2.1 to rank the possible alignments (3Di+AA), with two more alignment results using TM score and LDDT. Then three distance matrices are generated respectively from the three structural alignment results. Foldtree subsequently constructs three phylogenetic trees using the matrices with QuickTree (Howe et al., 2002) via the neighbour joining (NJ) method and roots the trees with the Minimal Ancestor Deviation (MAD) method (Tria et al., 2017).

To compare the structure-based NJ tree with a standard sequence alignment-based NJ tree, the complete sequence dataset was aligned in the programme Seaview V. 5.0.5 (Gouy et al., 2010) with Clustal Omaga (Sievers et al., 2011), using ten combined guided-tree and HMM iterations. The NJ tree was also calculated in SeaView (Gouy et al., 2010) using 1000 bootstrap replicates, observed distances, and ignoring all gap sites.

## Result and discussion

### Structural comparisons

AlphaFold uses predicted Local Distance Difference Test (pLDDT) scores to show the confidence of the prediction. It estimates per-residue C_α_ scores between predicted model and the true structure (Mariani et al., 2013, Jumper et al., 2021). The pLDDT value is in the range of 0-100, and over 90 is regarded as at the high accuracy level of an experimentally determined structure. Between 70 and 90 is good enough for backbone prediction, and below 70 needs careful interpretation whereas below 40 generally indicates low confidence in the prediction (Jumper et al., 2021, Terwilliger et al., 2024). Several studies have found that the pLDDT score is a good indicator for the flexibility of the region, with lower values indicating more dynamicity (Guo et al., 2022, Saldano et al., 2022, Ma et al., 2023). For a more intuitive visualisation of pLDDT score, all the predicted structures in figures follow pLDDT colouring scheme, i.e., regions over 90 are in blue, between 70 and 90 are in sky blue, between 50 to 70 in yellow, and below 50 in orange.

Predicted D1 structures achieved high average pLDDT scores overall (Figure 1), with important regions for structural integrity, such as membrane helices, reached pLDDT scores over 90. In contrast, sequence specific extensions at both termini and unique sequence insertions gave low scores below 40. The tail at the N-terminal surface alpha helix (Na in Figure 1C) showed a stiff increase of pLDDT from the amino acid (AA) positions 1 to 11. This tail is unstructured and not resolved in experimental determined structures, although in a few of the AF structures it appears to fold as an alpha helix (Figure 1A in D1^FRL^, G3, G1). From positions 11 to 21, a well conserved surface alpha helix, important in recognition and degradation by FtsH proteases (Komenda et al., 2007), had a high score (>90) in all predicted structures. At the C-terminus, the score reached 80 at H332, which is a coordinating ligand to Mn1 in the Mn_4_CaO_5_ cluster (Umena et al., 2011). After this position, relatively low pLDDT scores were observed ranging from 40-60. This region includes the remaining residues binding the cluster, including E333, H337, D342 and A344, where the CtpA protease cleaves the peptide bond during assembly of the complex and prior to photoactivation (Anbudurai et al., 1994, Satoh and Yamamoto, 2007).

**Figure 1.**
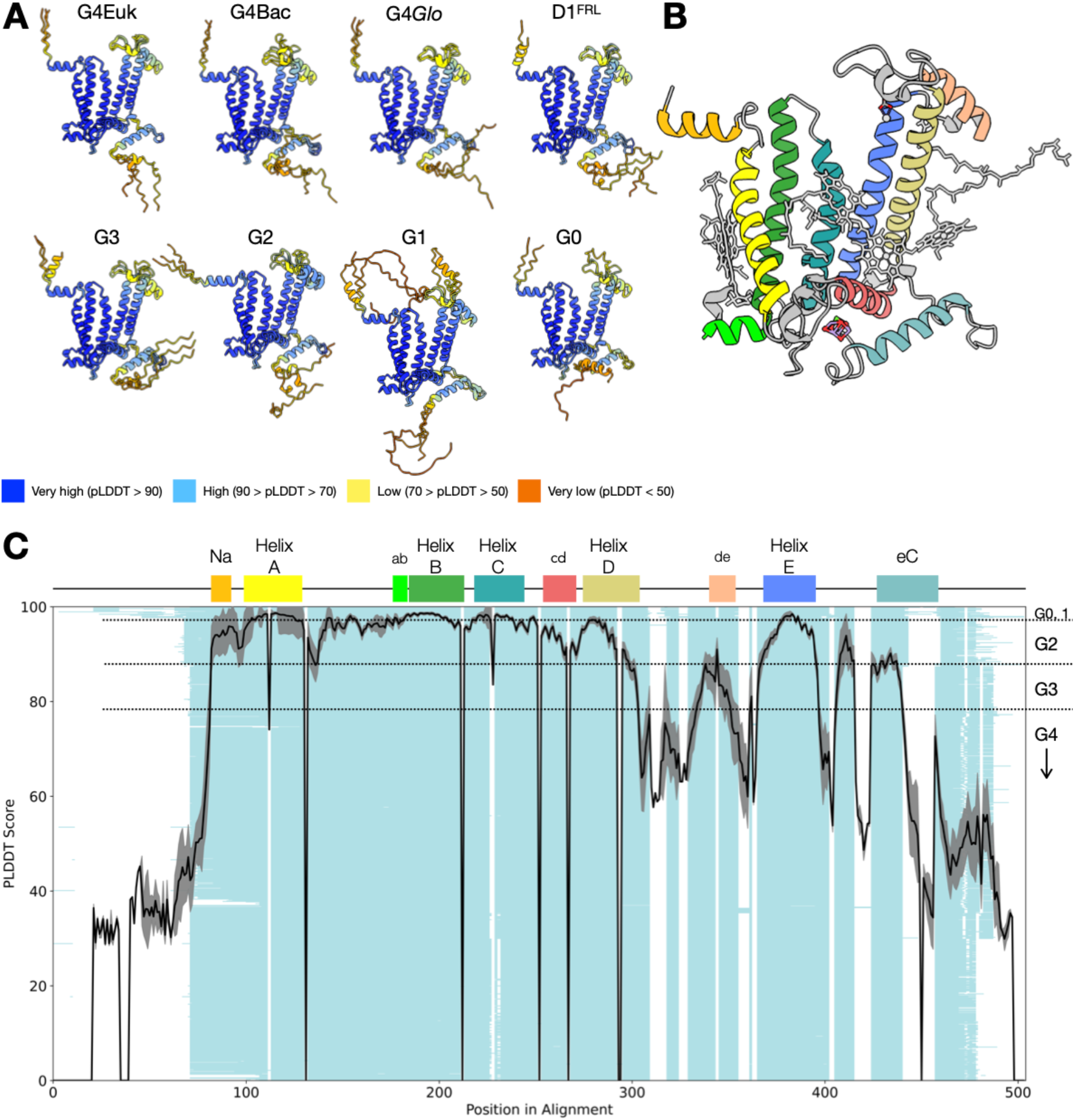
Representative structures from AlphaFold predictions and average pLDDT scores by amino acid position. **A** depicts the predicted AF structures from different D1 groups with pLDDT colouring scheme indicative of the pLDDT score by position. Every group shows an alignment of 3 representative structures except G0, which is comprised of two structures. **B** shows the recent CryoEM structure of the G4 D1 of *Synechocystis* sp. PCC 6803 (PDB ID 7N8O) as a reference. **C** shows the average pLDDT score by position (black line) overlaid on the sequence alignment of the same D1 dataset (light turquoise) to indicate corresponding amino acid positions. Standard deviation of pLDDT scores is shown in grey. The secondary structure annotation is depicted on top of the plot to illustrate the locational context in D1 and the colour scheme corresponds to that of panel **B**. Helix A to E denote transmembrane helices, Na denotes the N-terminal helix, eC denotes C-terminal helix, and ab, cd and de denote α helices in between the transmembrane helices. The D1 grouping in the sequence alignment is indicated with dotted lines on the right side of the plot.

These unstructured or flexible regions were also pronounced in the structural alignment analysis. Figure 2A shows the 23 representative AF structures from different D1 groups aligned to the reference structure. For example, predicted folding at the N-and C-terminus includes random coils and helices with different torsion angles to the N-terminal helix or C-terminal helix. Conversely, the common core in Figure 2C broadly agrees with the high pLDDT regions in Figure 1C and 2A. Notably, while loops connecting the A, B, C and D helices, i.e., AB, BC, and CD loops are included in the common core and have high pLDDT scores (>90), the DE loop, which includes the binding site of the mobile plastoquinone Q_B_ (the terminal electron acceptor of PSII) and is important for the proper assembly of PSII (Mulo et al., 1997, Ferreira et al., 2004), is only partially conserved. It suggests that the DE loop might be a variable region, allowing functional diversification. The DE loop also shows varying pLDDT score, reaching 80-85 in the regions between T228-S232, T248-L258 and at S264 and F265, while the rest of the region has relatively low scores of 60-70.

**Figure 2:**
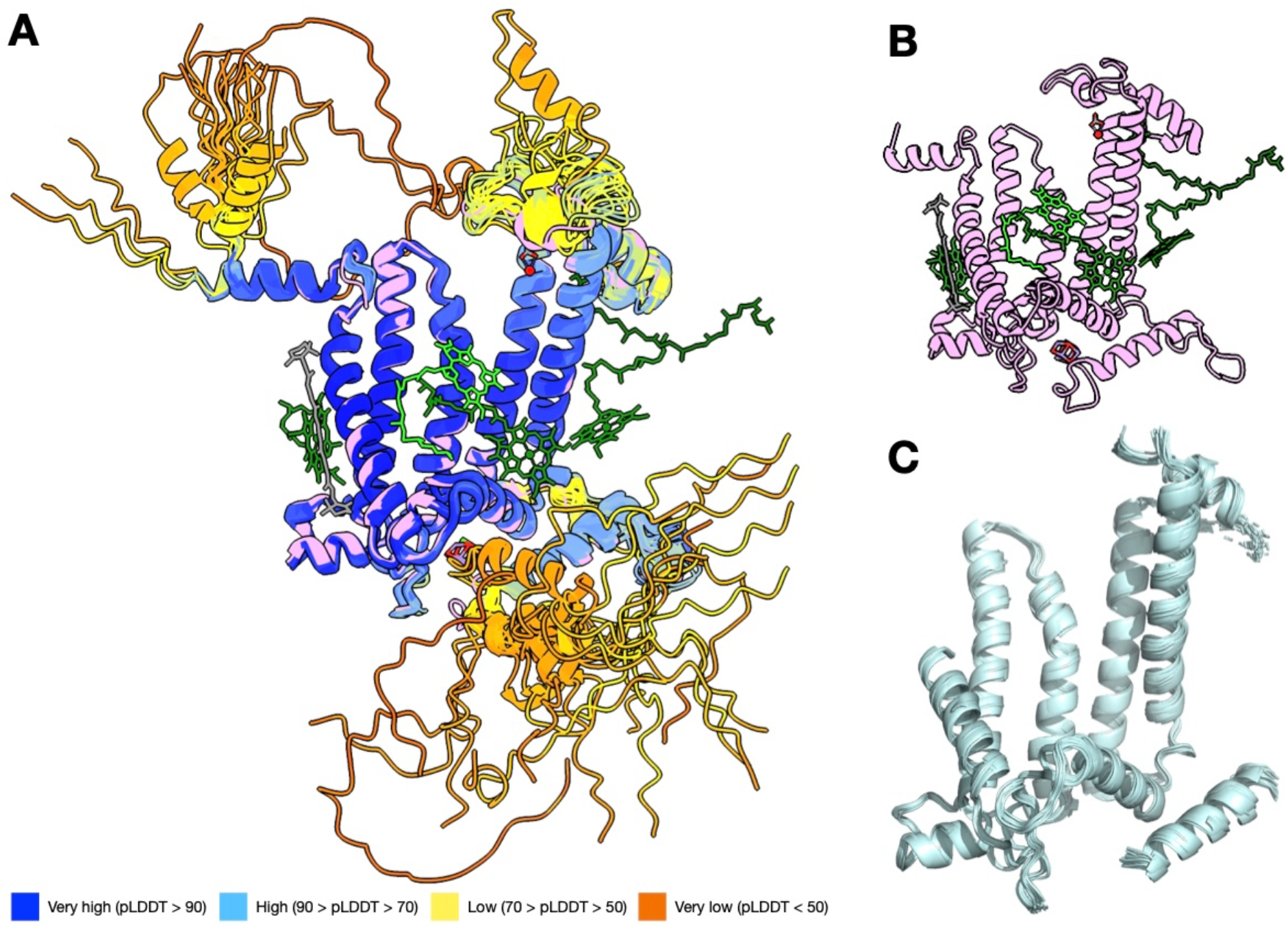
Structural alignment analysis with mTM-Align comparing global similarity of the full-length proteins. **A** shows the global alignment of 23 representative D1 AF structures from different D1 groups in pLDDT colouring, aligned to the reference G4 D1 (PDB ID:7N8O). The reference structure is depicted alone in **B** for comparison, and **C** depicts the common core illustrating the structurally conserved regions.

At the C-terminus, the common core ended at position H332 of the terminal eC helix. This is consistent with the diversification of the donor site of D1, particularly found in the atypical D1 groups (G0-G2), where several of the ligands to the cluster have been lost as they evolved to perform functions other than water oxidation (Oliver et al., 2023). In addition, we also noted that the N-terminus, including a part of helix A was missing in the calculated common core, despite the region retaining a high degree of sequence identity and high pLDDT scores (positions 12-35). This may be caused by the inclusion of a few partial sequences in the dataset and sequences that appeared to start at a different codon, which may represent annotation errors.

To get insight into the functional implications of the pLDDT variation in the DE loop region, we inspected the representative structures in further detail. Figure 3 highlights the diverse DE loop configurations predicted in the different groups of D1. The region between T248 and L258 is a main part of a hydrophobic cavity accommodating the Q_B_ (Ferreira et al., 2004), including H252, a key residue in the protonation of Q_B_ (Saito et al., 2013). The S264 and F265 make part of the Q_B_-binding site, providing two hydrogen bonds to the distal oxygen, and it is thought that S264 is also involved in one of the protonation routes (Saito et al., 2013). Both regions are structurally well conserved across different groups. Given their essential role in the binding, reduction and exchange of Q_B_, it is expected that the cavity and the coordinating residues would be subjected to negative selection pressure, thus preventing structural alteration. By contrast, the region between R225 and Q241 showed more diversity in its configuration with low average pLDDT scores except for T228-S232, the latter being a site of protein-protein interactions with CP47 and PsbL, while the low scoring residues appear exposed to the cytosol. The G0 and G1 forms are the most divergent in this regard, showing differences in length and secondary structure. Previous work had found that deletion of the R225-F239 region results in a strain still capable of photoautotrophic growth and oxygen evolution, but has impaired Q_B_ reduction (Mulo et al., 1997). Therefore, structural divergence could be permissive in this region (Pascual-Garcia et al., 2010) and it is plausible that mutations here could modulate electron or protonation rates at the acceptor side. Furthermore, recent structural work on intermediate PSII complexes during the assembly process has shown that the binding of the assembly factor, Psb28, triggers a significant conformational change of the region between residues L223-E244 into an extended beta hairpin structure (Xiao et al., 2021, Zabret et al., 2021). Our results are consistent with this region retaining a degree of flexibility as it coincides with the zone of low pLDDT scores, suggesting that mutations in this region could also modulate the rate of assembly by affecting beta hairpin folding and protein-protein interactions with Psb28. In the case of the most divergent forms of D1, such as G0 or G1, these interactions may be altogether abolished as the complex would not have the capacity to support water oxidation.

**Figure 3.**
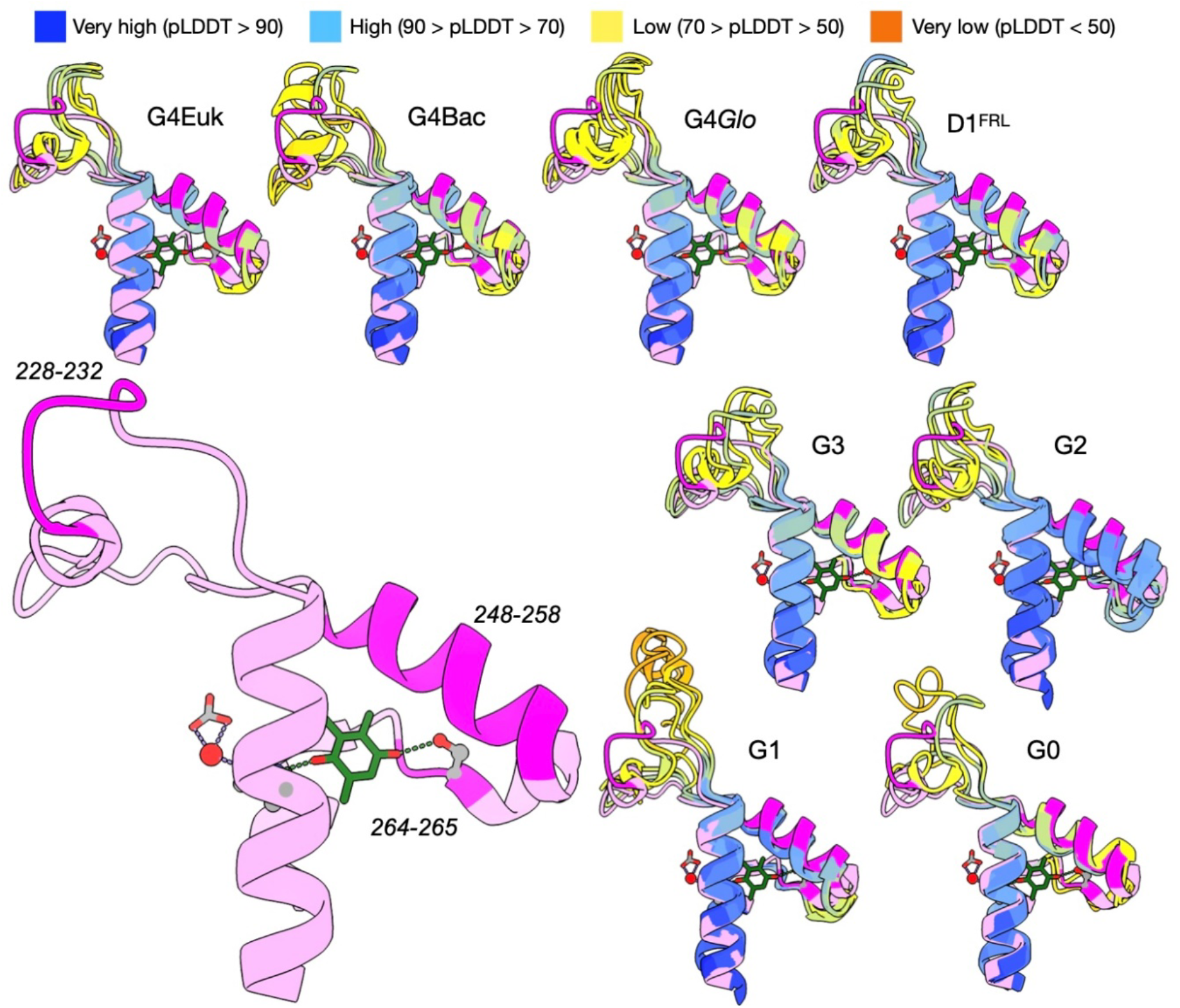
Diversity of DE loops in D1. The reference structure is shown in pink at the bottom left. Sections with high pLDDT scores are shown in magenta and their AA positions are indicated next to them. AF structures are aligned by group, and the reference structure is included in all the alignments for comparison. AF structures are depicted using the pLDDT colouring scheme. Q_B_ is shown in green and the lipidic tail is omitted for clarity. The non-heme iron is shown in red coordinated by a bicarbonate molecule in grey.

Notably, all the AF structures of the 23 representatives showed different configurations compared with the reference structure in terms of groove and 3D coordinates of the corresponding DE loop residues, even the standard G4 D1 proteins. In fact, the AlphaFold predicted PsbA2 protein from *Synechocystis* sp. PCC 6803, the reference D1 of this study, showed similar deviations in the DE loop region relative to the experimentally determined structure (Figure S2). It might indicate a conformational rearrangement of the region after initial folding, possibly facilitated by interactions with surrounding subunits (Figure S3), or a systematic deviation of the prediction from experimental structure due to the dynamic nature of the loop region. Taken together, these observations suggest that the DE loop might be comprised of two distinctive regions (R225-Q241 and T248-F265) diverging at different speeds, with the faster evolving regions potentially tuning the kinetics of Q_B_ reduction and the rates of assembly depending on environmental or physiological conditions and the slower evolving regions involved in the binding and protonation of Q_B_.

Figure 4 shows the structural diversity around the Mn_4_CaO_5_ cluster and the C-terminus. Most of the lumenal region of D1 up to position H332 at the C-terminus aligns well with the reference structure, including the two ligands of the Mn_4_CaO_5_ cluster D170 and E189. In the C-terminus, the loop between helix E and the C-terminal helix does not deviate from the reference structure, except for G1 D1, which shows a beta hairpin between residues 305-315, see also Figure S4. This loop is a site of protein-protein interactions in D1, providing a binding site to both subunits of the cyt *b*559, PsbV, and featuring protein-protein interactions with D2. The presence of a beta hairpin in this region could suggest either a disruption of protein-protein interactions with the other subunits in the “super-rogue PSII” involved in the synthesis of chlorophyll *f* (Trinugroho et al., 2020); or alternatively, new interactions with one or more novel subunits. Purification of PSII complexes from *Synechocystis* sp. PCC 6803 with a bound G1 D1 from *Chroococcidiopsis thermalis*, showed an absence of the extrinsic polypeptides and binding of the assembly factor Psb27 (Trinugroho et al., 2020).

**Figure 4.**
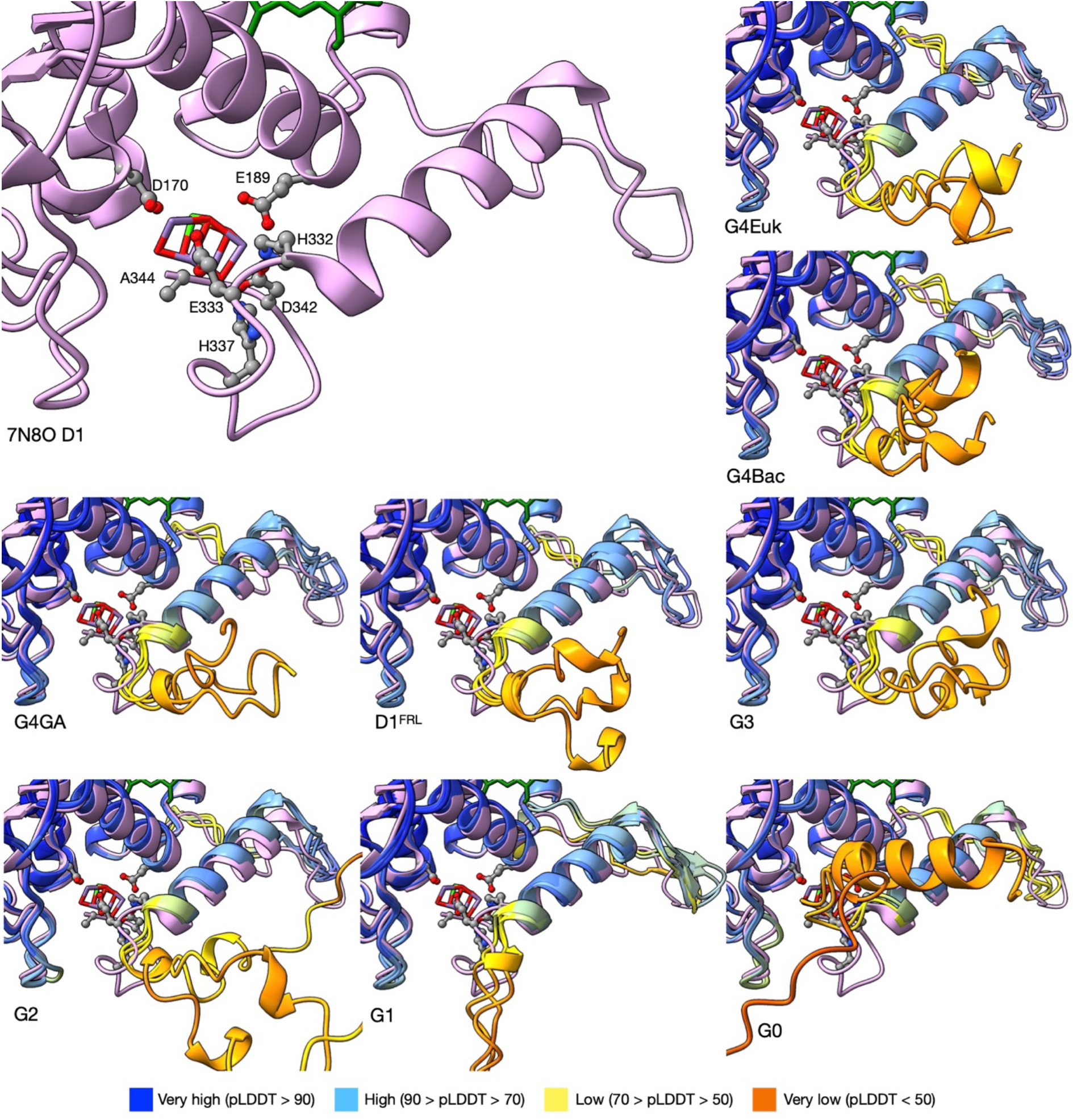
Details of the surrounding region of Mn_4_CaO_5_ cluster and C-terminus, aligned by group. The reference structure is shown in pink on the top left and the ligands for Mn_4_CaO_5_ cluster are labelled next to their atomic depiction. AF structures are aligned with the reference structure for comparison, using pLDDT colouring. Residues after A344 are hidden in G3 and G4 to account for the cleavage by CtpA protease.

The C-terminal alpha helix (eC) shows high pLDDT scores overall across D1 groups, but the score becomes low after position H332 and then very low after position N335, possibly suggesting increasing flexibility towards the end of the C-terminal loop. Indeed, an occasional short helix is also observed across the D1 groups in the very low pLDDT region, following no specific evolutionary pattern. This short helix is quite prominent in the G0 D1 found in the genome of *Gloeobacter morelensis*, being 12 AA in length, but this is absent in the G0 D1 of *G. kilaueensis*. The function of this D1 is still unknown, and whether this D1 is expressed and incorporated into PSII still remains to be determined.

In terms of the torsion angle of the C-terminal loop to the eC helix, all predicted structures protrude towards the opposite direction of the loop in the reference structure. That is, away from the Mn_4_CaO_5_ cluster. To check if the unfolded C-terminal loop of the AF structures agreed with experimental structures we compared them with the cryoEM structures of PSII intermediates during assembly with bound Ycf48 (Zhao et al., 2023) or Psb27 (Zabret et al., 2021), which feature alternative configurations of the C-terminal loop (Figure S5), but they showed no overlap. This positioning of the loop before cleavage could potentially represent an evolutionary adaptation to minimise the risk of partial photoactivation of the cluster during the assembly process before the complex is ready to operate safely, as all ligands are available in unprocessed D1, except for A344’s carboxylic terminus.

### Structural phylogenetics

Next, to examine how the evolutionary information transferred from sequence to structure, Foldtree (Moi et al., 2023) was used to generate a phylogenetic tree from the AF structures (Figure 5). The tree topology follows a similar branching pattern to the tree reported in previous studies (Murray, 2012, Cardona et al., 2015, Sheridan et al., 2020), which was inferred used Maximum Likelihood (Figure 5A). The atypical and very divergent G0 sequences represent the first branching point, with D2 as the outgroup, then followed by G1 to G4. G3, which contains sequences known to be expressed under low-oxygen conditions, also referred to as D1’, no longer appear as a monophyletic group, but it is split into three separate clades, suggesting that not all D1’ may be functionally equivalent, but reproduces what is observed in the sequence-only NJ tree (Figure 5C).

**Figure 5.**
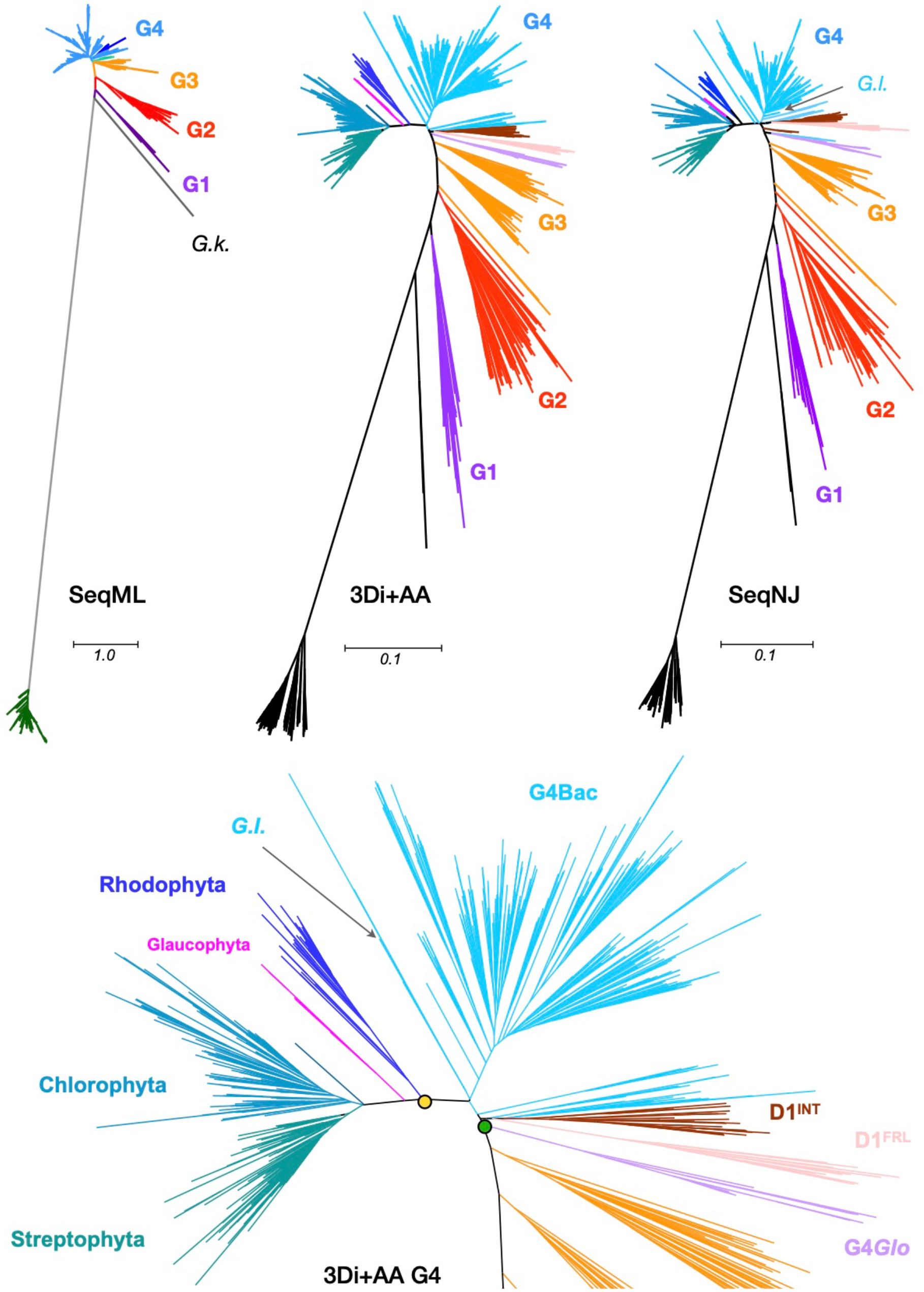
Phylogenetic trees of D1 proteins constructed from amino acid sequences and AF D1 structures. For comparison, **SeqML** shows the Maximum Likelihood tree of cyanobacterial D1 by Cardona et al. (2015), with *G*.*k*. indicating G0 from *Gloeobacter kilaueensis*. **3Di+AA** shows the phylogenetic tree constructed with AF D1 structures by Foldtree (Moi et al., 2023). **SeqNJ** shows the neighbour joining tree based on the D1 sequence dataset used in this study. Scale bars represent AA substitutions per site. **3Di+AA G4** shows the G4 D1 topology of the structural tree in more detail, with D1^INT^ indicating “intermediate D1” as described in Sheridan et al. (2020). *G*.*I*. in **SeqNJ** and **3Di+AA G4** indicates the only D1 sequence found in the genome of *Gloeomargarita lithophora*, which is considered as the closest living cyanobacterium to the ancestor of the chloroplast. Green and yellow circle indicate the most recent common ancestor (MRCA) of cyanobacteria and primary plastid, respectively. Glaucophyta are shown in magenta.

The diversification of D1 within photosynthetic eukaryotes in the structural phylogeny (Figure 5B) follows those observed before, with the red algae as the earliest branch, and with Glaucophytes and the green lineage sharing a common ancestor (Sanchez-Baracaldo et al., 2017, Bowles et al., 2024), supporting the monophyletic origin of the primary plastid. Furthermore, the structural tree located *Gloeomargarita lithophora* in the close vicinity of the Most Recent Common Ancestor (MRCA) of the primary plastids, which is also in agreement with earlier phylogenomic analyses (Sanchez-Baracaldo et al., 2017, Ponce-Toledo et al., 2017, Moore et al., 2019). In contrast, the sequence-based neighbour joining (NJ) tree (Figure 5B) was not able to recover the same topology, showing Glaucophytes as the earliest branch of Archaeplastida mixed with other algae, and placed *Gloeomargarita lithophora*, somewhere else in the tree.

A notable difference of the structural tree compared with previous sequence-based phylogenetic studies, as well as the sequence-only NJ tree (Figure 5C), concerns the topology of the G4 sequences (Figure 5D), which is the standard D1 used for water-splitting under most conditions. Given that all cyanobacteria retain at least one, but often several G4 D1, and assuming that G4 D1 has been inherited mostly vertically across cyanobacteria, one could argue that the branching point of G4 D1 in Gloeobacterales, indicates or approximates the MRCA of cyanobacteria. In previous studies (Sheridan et al., 2020, Gisriel et al., 2022a), the D1^FRL^ was located in between G3 and G4, thereby suggesting that the gene duplication leading to its origin predated the MRCA of cyanobacteria. This branching pattern, taken together with the early duplication of the D1 forms leading to the emergence of G1 (the chlorophyll *f* synthase), would open the possibility that an acclimation response to far-red light, involving the synthesis of at least chlorophyll *f*, originated before the radiation of the extant groups of cyanobacteria. The current result suggests a simpler scenario, that D1^FRL^ is a G4 D1, which specialised early during the diversification of cyanobacteria. This reduces the number of D1 copies the MRCA of cyanobacteria may have had. This scenario is also in line with the evolutionary history of other FRL PSII subunits in the FaRLiP gene cluster, which were found to have diverged after the MRCA of cyanobacteria in their respective phylogenies (Gisriel et al., 2022a). The same logic can be applied to D1^INT^, defined by Sheridan et al. (2020) as a subgroup of intermediate D1 between G3 and G4 and representing a D1 variant with an unknown function. In the structural phylogeny, D1^INT^ was also found to branch within G4, just immediately after the divergence of G4*Glo*. Furthermore, the structural phylogeny supported the close relationship between D1^FRL^ and D1^INT^, as seen in Sheridan et al. (2020), suggesting that these two may have specialised from the same ancestral form of G4 D1.

### Concluding remarks

The AlphaFold prediction revealed a higher structural diversity in D1 proteins than is currently observed experimentally, suggesting the possibility of novel functional specialisations. In addition, we found that structure-enhanced and sequence-based trees were in reasonable agreement, encouraging the use of phylogenetic analysis with predicted structures as a powerful tool to guide further research on the origin and evolution of photosynthesis.

Nevertheless, as much as AlphaFold opened a new era for structural bioinformatics with its prediction accuracy, there is still room for improvement. For example, the predictions did not include any of the core cofactors bound by D1, although it was recently shown that AlphaFold2 predicted structures of G-coupled receptors resembled more closely those with bound ligands than those without (Karelina et al., 2023). Excitingly, improvements on those aspects are already on the way. The latest version of AlphaFold, AlphaFold3 (Abramson et al., 2024), can accurately predict biomolecular interactions such as protein-protein, protein-DNA, protein-ions or binding small molecules, including chlorophyll *a* and *b*, as well as bacteriochlorophyll *a* and *b*. There are significant developments in the field on better methods to leverage structural information into phylogenetics, such as calculating bootstrap values from structural information (Baltzis et al., 2025), constructing clustered dataset from PDB for structural phylogenetics (Malik et al., 2023), and a rapid search method for structural similarity in a clustered AlphaFold protein database (Barrio-Hernandez et al., 2023).

Structural phylogenetics allows distant relationships over the limit of detectable sequence homology to be found, as structural features evolve slower than sequences (Illergard et al., 2009, Malik et al., 2020). With the help of the explosive advances in the structural bioinformatics field, we envisage that AlphaFold-powered structural phylogenetics will be a promising strategy to illuminate the entire map of photosystem evolution, including fully assembled type I and II reaction centres from phototrophic prokaryotes to plants.

## Author contributions

T.D.K. designed the experiments; T.D.K. and D.P. carried out the experiments; T.D.K. and T.C. analysed the data and wrote the manuscript; T.D.K, D.P. and T.C. edited the manuscript; J.M.W. edited the manuscript and provided scientific input. T.C. secured funding for the project and provided supervision.

## Data availability statement

The data that supports the findings of this study are available in the supplementary material of this article, Predicted AF structures, structural alignment analysis and phylogenetic trees are available at https://doi.org/10.5281/zenodo.14981431.

## Acknowledgement

The generous support of a UKRI Future Leaders Fellowship is gratefully acknowledged (MR/T017546/1, MR/T017546/2, and MR/Y011635/1 to T.D.K. and T.C.). D.P. is supported by an Imperial College London EPSRC DTP studentship.

