## Supplementary material for "Exploring the structural diversity and evolution of the D1 subunit of photosystem II using AlphaFold and Foldtree"

Table S1. Full list of 24 representative structures

| Group | Species origin |
|---|---|
| G0 | *Gloeobacter morelensis* |
| G0 | *Gloeobacter kilaueensis* |
| G1 | *Chroococcidiopsis* sp. CCALA 051 |
| G1 | *Synechococcus* sp. PCC 7335 |
| G1 | *Fischerella thermalis* |
| G2 | *Anabaena* sp. PCC 7108 |
| G2 | *Synechococcus* sp. PCC 7336 |
| G2 | *Nostoc* sp. ATCC 53789 |
| G3 | *Synechococcus* sp. PCC 7310 |
| G3 | *Pseudanabaena* sp. PCC 6802 |
| G3 | *Leptolyngbya* sp. PCC 7375 |
| $D1^{FRL}$ | *Chroococcidiopsis* sp. CCALA 051 |
| $D1^{FRL}$ | *Chlorogloeopsis fritschii* |
| $D1^{FRL}$ | *Synechococcus* sp. PCC 7335 |
| G4*Glo* | *Gloeobacter morelensis* |
| G4*Glo* | *Gloeobacter kilaueensis* |
| G4*Glo* | *Candidatus* Cyanoaurora vandensis |
| G4Bac | *Fischerella* sp. PCC 9605 |
| G4Bac | *Calothrix* sp. 336/3 |
| G4Bac | *Synechococcus* sp. PCC 7002 |
| G4Euk | *Flintiella sanguinaria* |
| G4euk | *Pharus lappulaceus* YP 008 |
| G4euk | *Cannabis sativa* |
| Reference | *Synechocystis* sp. PCC 6803 (G4) |

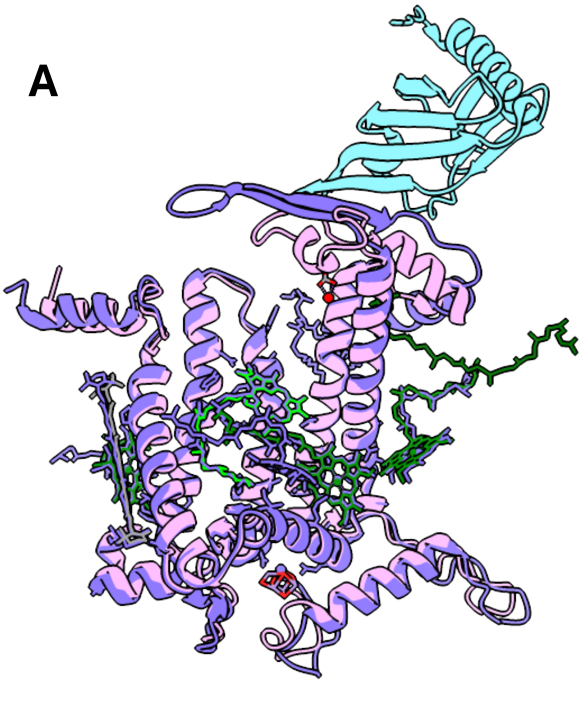


**Figure S1**. Experimentally determined structure of Psb28 bound D1 protein in an intermediate Photosystem II and the DE loop region in detail. The structure was determined by cryo-EM using *Thermosynechococcus vestitus* BP-1,PDB ID: 7NHP (Zabret et al., 2021). D1 is shown in purple and Psb28 in cyan. The structure is aligned to a reference structure: the D1 (PsbA2) subunit of *Synechocystis* sp. PCC 6803, PDB ID:7N8O (Gisriel et al., 2022) in pink.

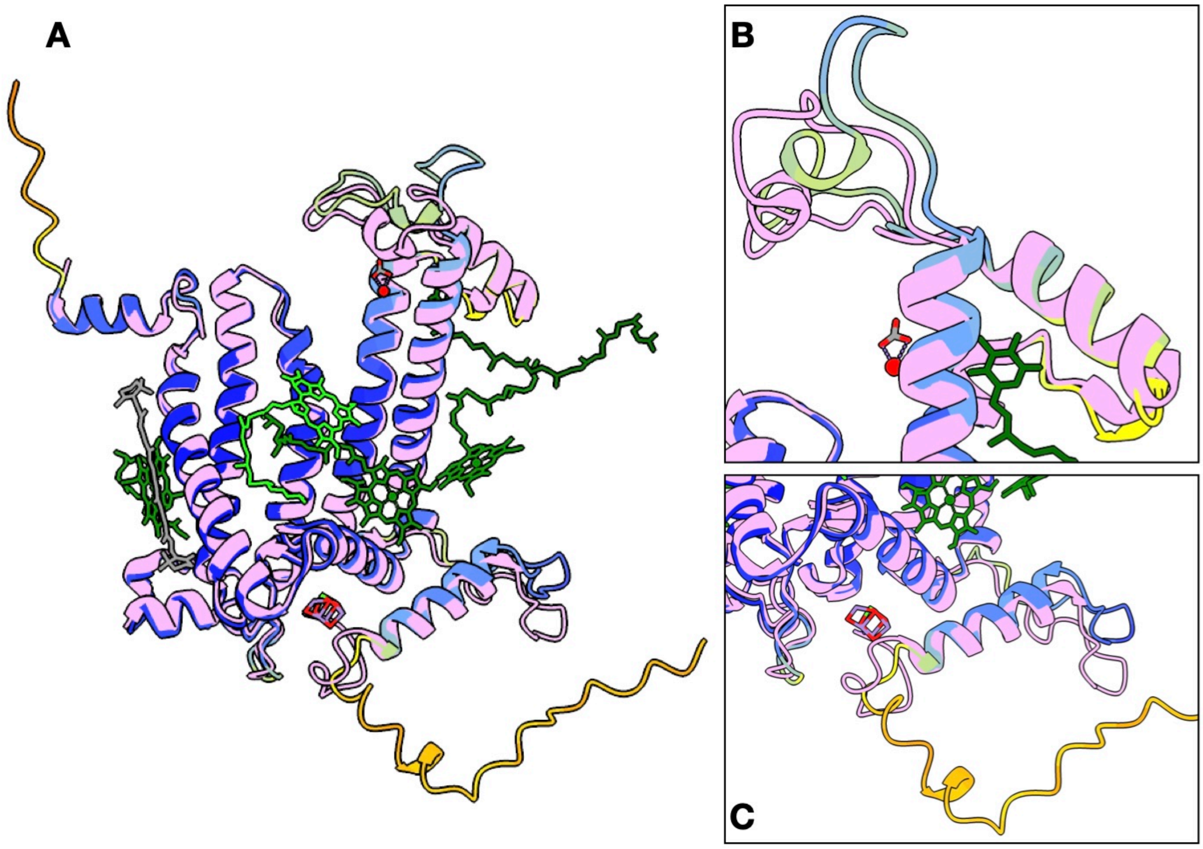


**Figure S2**. Comparison between experimentally determined and AlphaFold-predicted D1 protein. The experimentally determined structure (cryo-EM) of D1 in *Synechocystis* sp. PCC 6803 is shown in pink, and the AlphaFold prediction of the protein is depicted in pLDDT colouring. **A** shows the global alignment of the two structures; **B** highlights the DE loop region in detail and the lumenal region and C-terminus are shown in panel **C**.

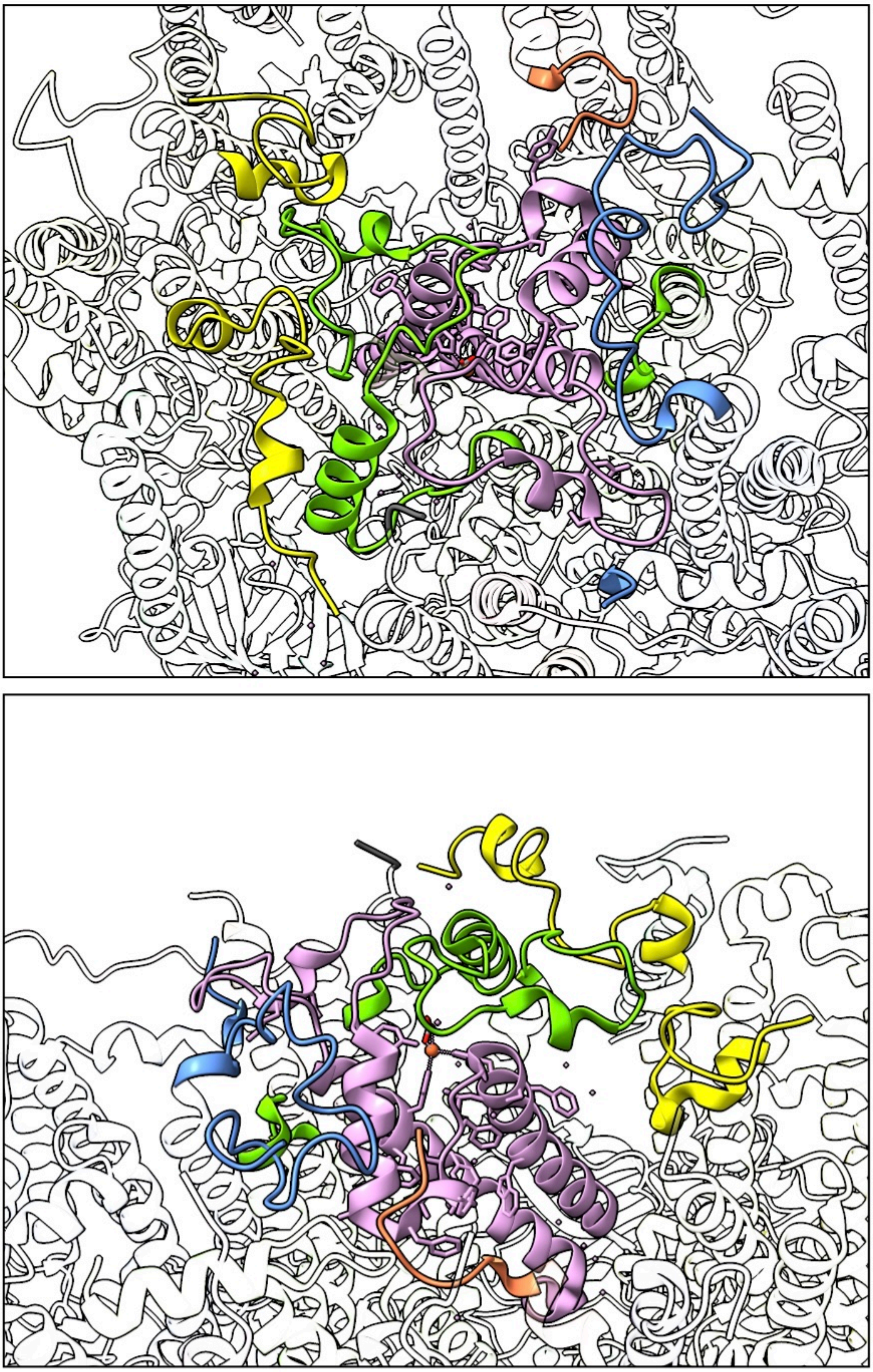

**Figure S3**. Surroundings of the DE loop in detail (PDB 7N8O). D1 is depicted in pink, CP43 in yellow, D2 in green and CP47 in blue. PsbE is depicted in orange. Top panel shows a view from the top of the enzyme (stromal side), while the bottom side has been rotated to show a different perspective on the site.

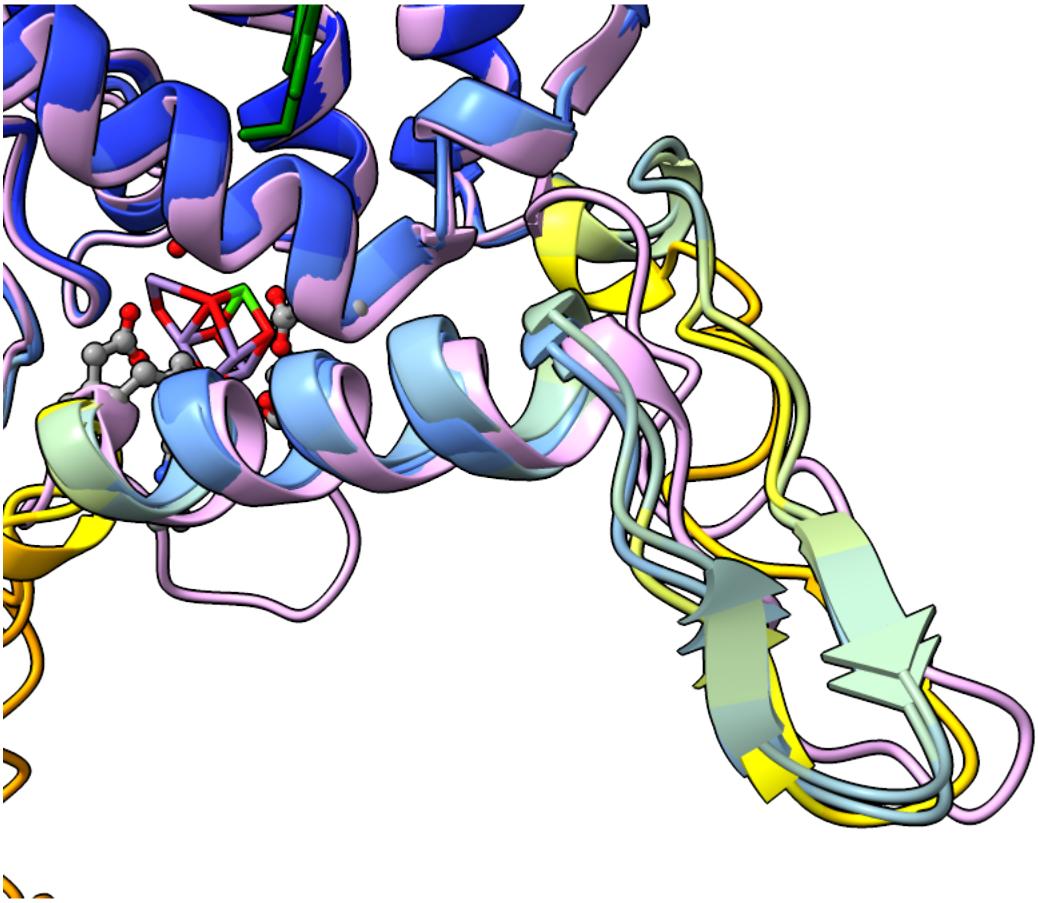

**Figure S4**. Close up of the beta-hairpin found in the C-terminal region of the G1 D1 structures. The reference structure is depicted in pink, and the other three G1 AF structures are in pLDDT colouring.

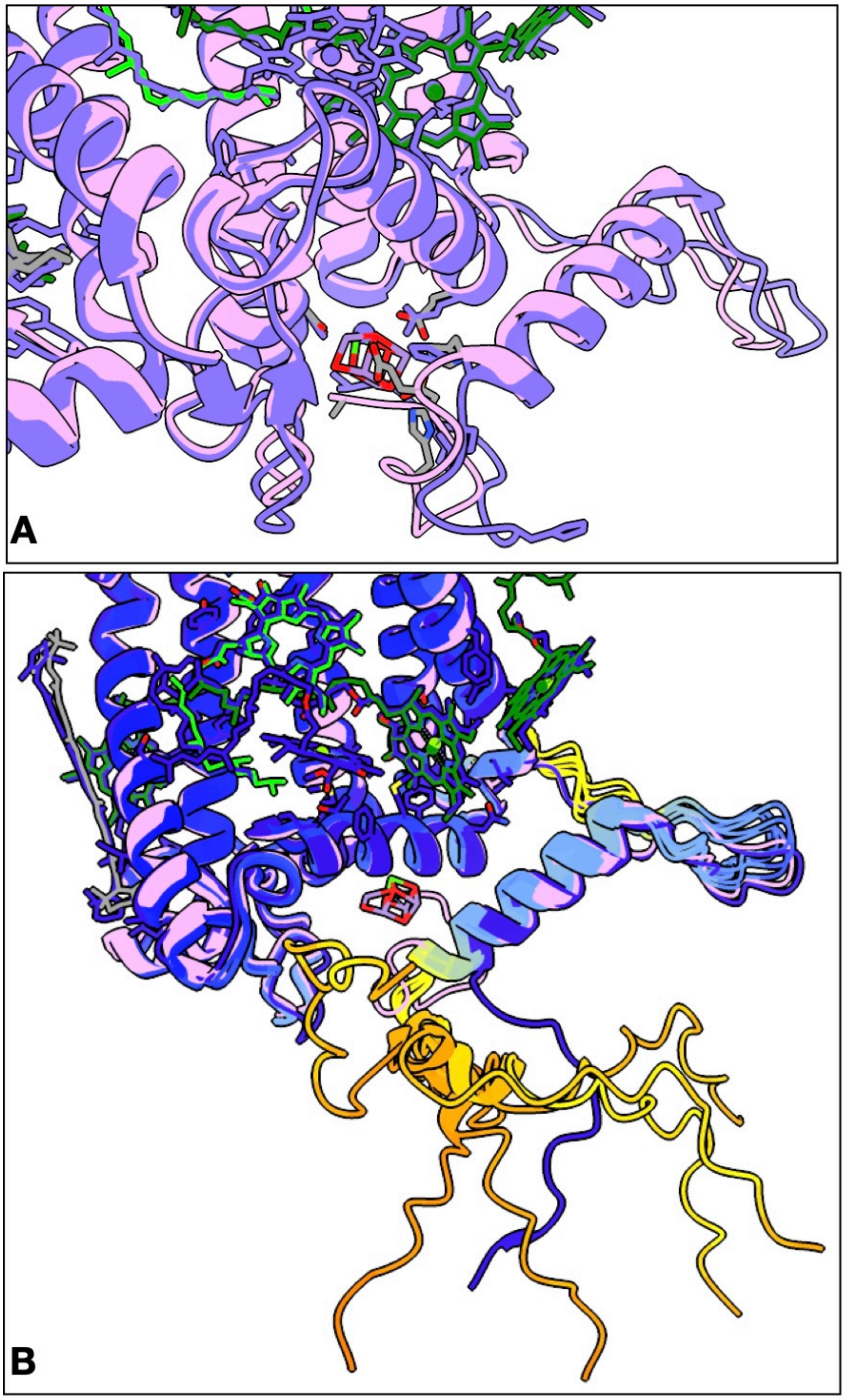


**Figure S5**. Comparison of the C-terminus of D1 from different PSII complexes during intermediate steps of assembly. **A** shows the C-terminus of D1 from an intermediate PSII with bound assembly factors Psb27, Psb28 and Psb34 reported by Zabret et al. (2021), PDB ID: 7NHP. The assembly factors are not shown for clarity. The reference structure is coloured pink. **B** shows the C-terminus from a PSII complex with bound Ycf48, instead of the extrinsic polypeptides PsbO, V, and U as reported by Zhao et al. (2023), PDB ID: 8AM5. This is shown in deep blue and the reference structure is in pink. G4Euk and G4Bac structures are shown in pLDDT colouring for comparison.
